# Inhibition of serotonin biosynthesis in neuroendocrine neoplasm suppresses tumor growth *in vivo*

**DOI:** 10.1101/2023.04.07.536013

**Authors:** Dane H. Tow, Catherine G. Tran, Luis C. Borbon, Maclain Ridder, Guiying Li, Courtney A. Kaemmer, Ellen Abusada, Aswanth Harish Mahalingam, Anguraj Sadanandam, Chandrikha Chandrasekaran, Joseph Dillon, Douglas R. Spitz, Dawn E. Quelle, Carlos H.F. Chan, Andrew Bellizzi, James R. Howe, Po Hien Ear

## Abstract

Small bowel neuroendocrine tumors (SBNETs) originate from enterochromaffin cells in the intestine which synthesize and secrete serotonin. SBNETs express high levels of tryptophan hydroxylase 1 (Tph1), a key enzyme in serotonin biosynthesis. Patients with high serotonin level may develop carcinoid syndrome, which can be treated with somatostatin analogues and the Tph1 inhibitor telotristat ethyl in severe cases. Although the active drug telotristat can efficiently reduce serotonin levels, its effect on tumor growth is unclear. This study determined the effect of serotonin inhibition on tumor cell growth *in vitro* and *in vivo*. The levels of Tph1 in various neuroendocrine neoplasms (NENs) were determined and the biological effects of Tph1 inhibition *in vitro* and *in vivo* using genetic and pharmacologic approaches was tested. Gene and protein expression analyses were performed on patient tumors and cancer cell lines. shRNAs targeting *TPH1* were used to create stable knockdown in BON cells. Control and knockdown lines were assessed for their growth rates *in vitro* and *in vivo*, angiogenesis potential, serotonin levels, endothelial cell tube formation, tumor weight, and tumor vascularity. *TPH1* is highly expressed in SBNETs and many cancer types. *TPH1* knockdown cells and telotristat treated cells showed similar growth rates as control cells *in vitro*. However, *TPH1* knockdown cells formed smaller tumors *in vivo* and tumors were less vascularized. Although Tph1 inhibition with telotristat showed no effect on tumor cell growth *in vitro*, Tph1 inhibition reduced tumor formation *in vivo*. Serotonin inhibition in combination with other therapies is a promising new avenue for targeting metabolic vulnerabilities in NENs.

## Introduction

Serotonin, also known as 5-hydroxytryptamine (5-HT), is a small metabolite synthesized from tryptophan and functions both as a neurotransmitter and a hormone(Bader, 2020). In the central nervous system, the key enzyme that produces serotonin is tryptophan hydroxylase 2 (Tph2). Elsewhere in the body, tryptophan hydroxylase 1 (Tph1) is the main enzyme that regulates serotonin synthesis. *TPH1* is abundantly expressed in the pituitary, small bowel, and stomach (Park et al., 2021). Many tumors produce and secrete serotonin, which has autocrine effects through binding to 5-HT2B and 5-HT7 receptors on the tumor cell surface, which stimulates cell proliferation signaling cascades (Gautam et al., 2016, Jiang et al., 2017, Asada et al., 2009). In addition, serotonin can stimulate the biosynthesis of pro-fibrotic factors on the 5-HT1B receptor on cancer associated fibroblasts (CAFs) (Seuwen et al., 1988) in a paracrine fashion, and promotes angiogenesis in several cancers (Nocito et al., 2008, Nemecek et al., 1986, Asada et al., 2009). Serotonin has also been shown to dampen T-cell function to promote tumor growth in a colon adenocarcinoma mouse model (Schneider et al., 2021, Liu et al., 2021).

Neuroendocrine neoplasms (NENs) are rare cancers, which include well-differentiated neuroendocrine tumors (NETs) and poorly differentiated neuroendocrine carcinomas (NECs) (Rindi et al., 2018). Small bowel NETs (SBNETs) are well-known for their ability to secrete high levels of serotonin, which is also used as a diagnostic marker for this cancer (Bellizzi, 2020b, Bellizzi, 2020a). Patients with metastatic SBNETs often develop carcinoid syndrome due to the high level of serotonin released by cancer cells. Patients with severe diarrhea which cannot be managed with somatostatin analogues (SSAs) alone may be treated with the prodrug telotristat ethyl (Kulke et al., 2014, Dillon and Chandrasekharan, 2018). The active drug telotristat isa specific competitive inhibitor of Tph1 (Camilleri, 2011). Some pancreatic NETs and NECs also secrete serotonin, but those tumors are less commonly associated with carcinoid syndrome. The role of serotonin in NEN tumorigenesis remains underexplored. A study by Herrera-Martinez et al. showed no effect of serotonin on tumor cell growth in cell culture experiments(Herrera-Martinez et al., 2020) while a clinical study Morse et al. demonstrated that serotonin inhibition impairs tumor growth in patients (Morse et al., 2020) which is also supported by a case report with similar outcome (Molina-Cerrillo et al., 2016). Due to the limited number of cell lines and mouse models of NENs, only a few studies have reported on the role of serotonin regulation in NENs. BON cells are a well-established NEN cell line that reliable produces serotonin (Evers et al., 1991, Evers et al., 1994) and can be established as a xenograft mouse model that develops carcinoid syndrome (Musunuru et al., 2005, Jackson et al., 2009).

In this study, the effect of Tph1 expression, the rate limiting enzyme in serotonin biosynthesis, in NENs and other cancers was investigated. We confirm high levels of Tph1 expression in SBNETs and several lung NECs. In addition, genetic and pharmacological modulation of serotonin was used to understand its role in NEN tumor progression *in vitro* and *in vivo*. shRNA knockdown clones of *TPH1* and inhibition of Tph1 using telotristat in BON cells caused no effect on tumor cell growth *in vitro*. However, BON cells expressing *TPH1* shRNA or treated with telotristat resulted in decreased tumor growth and tumor vascularization *in vivo* as compared to their respective control groups.

## Materials and methods

### Cell culture and generation of *TPH1* knockdown cell lines

BON cells were gifted by Drs. M.B. Evers and C.M. Townsend and maintained at 37°C with 5% CO_2_ and cultured in DMEM/F12 medium supplemented with 10% Fetal Bovine Serum, 100 Units/mL penicillin + 100 μg/mL streptomycin, and 2 mM L-glutamine (Gibco; #26140079, #15140122, and #25030081). BON shControl and shTPH1 cell lines were generated using lentiviral transduction system with shRNA expression plasmids from Applied Biological Materials Inc: pLenti-Scrambled siRNA Control (# LV015-G), pLenti-TPH1_a (#i025101a), and pLenti-TPH1_c (# i025101c). Stable clones were selected using 2.5 μg/mL puromycin (AdipoGen; #AGCN20078). QGP-1 cells were purchased from the Japanese Collection of Research Bioresources (# JCRB0183). Lung, colon, and pancreas cancer cell lines (DMS53, DMS273, H727, H69, H69AR, H720, Colo205, SW1990, BxCP-3, PANC-1, and CFPAC-1) were purchased from the American Type Culture Collection. Cholangiocarcinoma cell lines (HuCC-T1 and EGl-1) were gifts from Dr. L.R. Roberts.

### Cell viability and growth assays

BON cells were treated with increasing concentration of telotristat (AdooQ; #A11785) or vehicle control for 3 days and assessed for cell viability using resazurin dye (MedChemExpress, #HY-111391). For cell growth assays, BON shControl, shTPH1_#1, and shTPH1_#2 cell lines were seeded at a density of 10,000 cells per well in 24-well plates. Cells from 4 different wells from each cell line were collected and counted with a hemacytometer on the respective days. Culture media were changed every 2 days for cells in the remaining wells. Error bars represent standard deviation (SD).

### Tube formation assays

Human umbilical cord vein endothelial cells (HUVECs, purchased from ATCC; #PCS-100-010) were cultured in endothelial cell growth medium (Cell Applications; #211). For tube formation assays, HUVECs were seeded at a density of 2,000 cells/well in 96-well plates and cultured in conditioned media from BON shControl, shTPH1#1, or shTPH1#2 cell lines with and without 1 μM serotonin supplementation (Sigma; #14927). Bright field images were take using an Olympus microscope (CKX53). HUVEC tube length was quantified by measuring the length cells continuously connected to each other using ImageJ 1.53a (National Institutes of Health). Statistical analyses of tube length from n = 3 experiments were performed using T-tests in GraphPad Prism Version 8.42. P <0.01 was depicted with **. P <0.0001 was depicted with ****. Error bars represent SD.

### Xenograft models

All animal experiments were approved in our animal protocol #2051771. Subcutaneous xenograft tumor models were generated by implanting 1 million BON cells in the flank of NSG mice (Jackson Laboratory; Stock #005557). Tumor volume was measured twice per week after tumor cell injection. After 10 days, mice were randomized into vehicle control and telotristat treatment groups (n = 4 mice per group). 30 mg/kg telotristat (AdooQ; #A11785) or an equal volume of vehicle control was administered to mice via intraperitoneal (IP) injections 5 days per week for a period of 17 days. Mice were euthanized at the end of the experiment and tumor samples were flash frozen in liquid nitrogen and fixed in formalin for further analyses. Statistical analyses of tumor weight were performed using T-tests in GraphPad Prism Version 8.42. P < 0.05 was depicted with *. P <0.01 was depicted with **. P <0.001 was depicted with ***. Error bars represent standard deviation.

Liver metastasis xenograft models were generated by injecting 1 million BON cells stably expressing the firefly luciferase gene (Kaemmer et al., 2021) in the spleen of an NSG mouse during surgery. Mice were given meloxicam daily for 7 days post-operation and monitored for signs of distress. On day 7 post-operation, mice were IP injected with 200 μL of 15 mg/mL D-luciferin salt (Gold Bio; # LUCK-100), incubated for 5 minutes, and imaged using the AMI HTX imager (Spectral Instruments Imaging, AZ) while anesthetized with 2.5% isoflurane. Bioluminescent signal was reported as photon flux corresponding to the number of photons per second that leave a square centimeter of tissue and radiate into a solid angle of one steradian (photons/sec/cm^2^/sr). Following baseline imaging, mice were randomized into vehicle control and telotristat treatment groups (n = 5 and n = 4 mice per group) and drug treatments were given 30 mg/kg telotristat 5 days per week for 14 days. On day 21 post-operation, bioluminescence tumor imaging was performed following the same protocol as the initial imaging. Mice were euthanized at the end of the experiment and tumor samples were flash frozen in liquid nitrogen or fixed in formalin for further analyses.

### Immunofluorescence staining

Patient tumors and mouse xenografts were embedded in Tissue-Tek OCT compound (Sakura; #4583) and frozen on dry ice prior to cryostat sectioning (∼4 μm) onto glass slides. Samples were fixed with 4% paraformaldehyde for 10 minutes and stained with anti-Tph1 (Sigma; #T0678) or anti-CD31 (BD; #550274) primary antibodies at 1:400 dilution overnight. Sections were then stained with anti-mouse FITC or anti-rat FITC (Jackson ImmunoResearch; #115-095-062 and #711-095-152) secondary antibodies at 1:500 dilution for 2 hours at room temperature. Fluorescent images were captured on a CKX53 microscope using cellSens software (Olympus) with a 300 ms exposure.

### Immunohistochemistry staining

Patient tumor samples were fixed in formalin and embedded in OCT for sectioning. Tissue sections were deparaffinized, rehydrated, and then heat-induced epitope retrieval was performed. Slides were stained with hematoxylin and eosin (H&E) and for Tph1 with an anti-Tph1 primary antibody (Sigma; #T0678). Detection was performed using the polymer-based Envision FLEX+ detection system (Agilent).

### Western blotting

BON shControl and shTPH1 cells were lysed in LDS Sample Buffer (Invitrogen; #NP0008) supplemented with 50 mM dithiothreitol. Whole cell protein lysates were briefly boiled, separated on a 4-12% polyacrylamide gel (Invitrogen; #NP0323BOX), and transferred to a PVDF membrane. Blots were incubated with anti-Tph1 R145 primary antibody (Sino Biological; #11931-R145) at a 1:1000 dilution followed by HRP-conjugated anti-rabbit secondary antibody (Jackson ImmunoResearch; #111-035-144) at a 1:3000 dilution. Chemiluminescent detection was performed using ECL substrate (BioRad; #1705060).

### Quantitative reverse transcription PCR

Surgically resected tissue and sbNET samples were collected from 5 patients under an IRB-approved protocol (IRB #199911057) between November 2019 and May 2021 and stored in RNAlater Stabilization Solution (Invitrogen; #AM7020). RNA was extracted from patient tissue samples and flash frozen mouse xenograft tumor samples (n = 4) using the RNeasy kit (Qiagen; #74104) and converted to cDNA using the qScript cDNA Synthesis kit (Quantabio; #95047). *TPH1*, *VEGF-A*, *VEGF-B*, and *VEGF-C* gene expression were assessed by qPCR on diluted cDNA samples using the oligos listed in Table 1 and PerfeCTa SYBR Green SuperMix (Quantabio; #95054). Expression level of *GAPDH* was used to normalize the expression of genes of interests across cDNA samples. Primer sequences were obtained from PrimerBank (https://pga.mgh.harvard.edu/primerbank/) and are listed on Table 1. All primers were purchased from Integrated DNA Technologies (IDT). Statistical analyses of gene expression changes were performed using T-tests in GraphPad Prism Version 8.42. P < 0.05 was depicted with *. P <0.01 was depicted with **. P <0.001 was depicted with ***. Error bars represent SD.

**Table 1.**
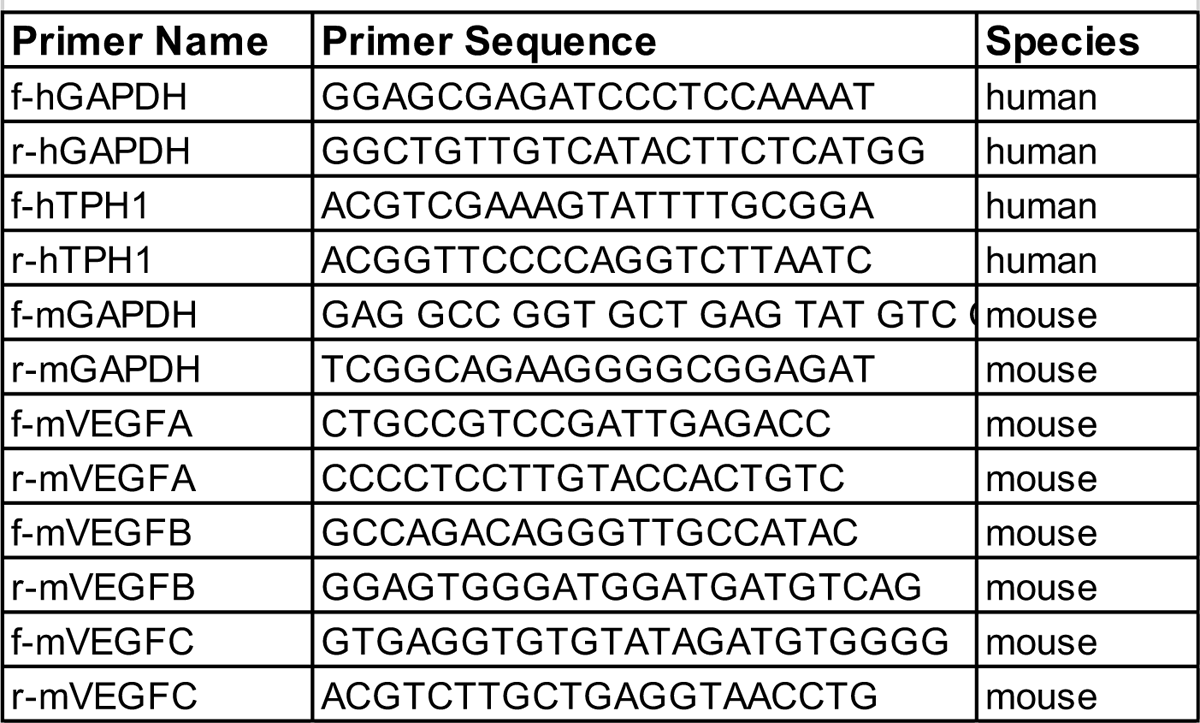
List of primers used in qPCR experiments.

### *TPH1* expression in TCGA database

The expression of the *TPH1* gene was analyzed across 33 TCGA cancer types using RNA-seq data and metadata obtained from the University of California Santa Cruz Xena browser (Goldman et al., 2020). The dataset consisted of 10,239 samples, including primary tumors (n=9,672), metastatic tumors (n=394), and primary blood-derived cancers (n=173). R version 4.2.2 and ComplexHeatmap 2.14.0 (Gu et al., 2016) and circlize 0.4.15 (Gu et al., 2014) packages were used to construct a heatmap.

## Results

### Tph1 is highly expressed in SBNETs

*TPH1* gene expression analysis of 5 sets of patient-matched normal small bowel tissue (SB), primary SBNETs, and SBNET liver metastases using quantitative reverse transcription PCR (RT-qPCR) with *TPH1*-specific primers (Table 1) demonstrated *TPH1* transcripts to be highly expressed in both primary SBNETs and metastases compared to adjacent normal SB tissue samples (Figure 1A). Tph1 protein expression in primary SBNET frozen tissue sections using immunofluorescent (IF) staining revealed Tph1 protein is highly expressed in tumor cells compared to the stroma (Figure 1B). Bright field imaging and DAPI nuclear staining (blue) are shown in the upper and lower panels, respectively. To better assess tumor morphology and cell-type specific expression, Tph1 protein expression was characterized by immunohistochemistry (IHC) staining on paraffin-embedded primary SBNET tissue sections (Figure 1C). Again, strong Tph1 expression was observed in the nested tumor cells and low expression in the surrounding stroma. H&E staining of an adjacent SBNET tissue section is shown in the Figure 1D for reference. Taken together, these data indicate Tph1 is highly and specifically expressed in SBNET cancer cells similarly to what was previously reported in a familial SBNET case (Sei et al., 2016).

**Figure 1.**
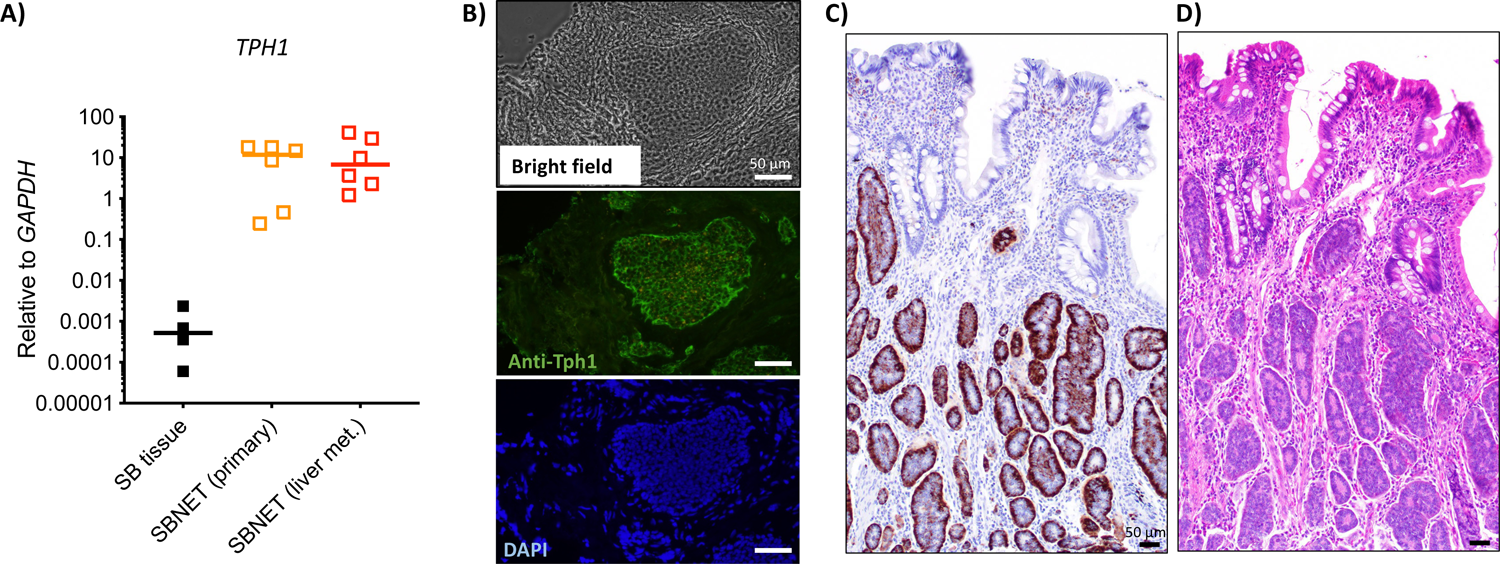
Expression of tryptophan hydroxylase 1 in SBNETs. (A) Expression levels of tryptophan hydroxylase 1 gene (*TPH1*) in primary SBNET and liver metastases (met.) compared to normal small bowel (SB) tissue. (B) Bright field image of an SBNET section. Tph1 protein expression detected by immunofluorescent staining using antibody specific against Tph1 (green) and nuclear staining using DAPI (blue). (C) Immunohistochemistry staining of an SBNET using specific antibody against Tph1 (brown). (D) H&E staining of an SBNET. Scale bars represent 50 μm.

### *TPH1* expression in other cancers

Analysis of *TPH1* expression levels in different cancer groups in The Cancer Genome Atlas (TCGA) database showed high *TPH1* expression in many cancer types such as acute myeloid leukemia, brain lower grade glioma, breast invasive carcinoma, esophageal carcinoma, glioblastoma multiforme, prostate adenocarcinoma, stomach adenocarcinoma, and uterine corpus endometrioid carcinoma (Figure 2A). The protein levels of Tph1 in BON cells (a pancreatic NEN line that expresses serotonin) were compared to QGP1 cells (a pancreatic NEN line that does not express serotonin) and other cancer cell lines (Figure 2B&C). Two pancreas cancer cell lines (PANC-1 and CFPAC-1) and 1 colon cancer cell line (SW1990) express Tph1 (Figure 2B). The lung cancer cell lines DMS53 (small cell carcinoma of the lung), H727 (non-small cell carcinoma), H69 (small cell lung carcinoma), H69AR (small cell lung carcinoma resistant to doxorubicin), DMS273 (small cell lung carcinoma) and H720 (atypical lung carcinoid) were analyzed for Tph1 expression. High Tph1 levels were found in in BON, H69AR, and DMS273 cells (Figure 2C).

**Figure 2.**
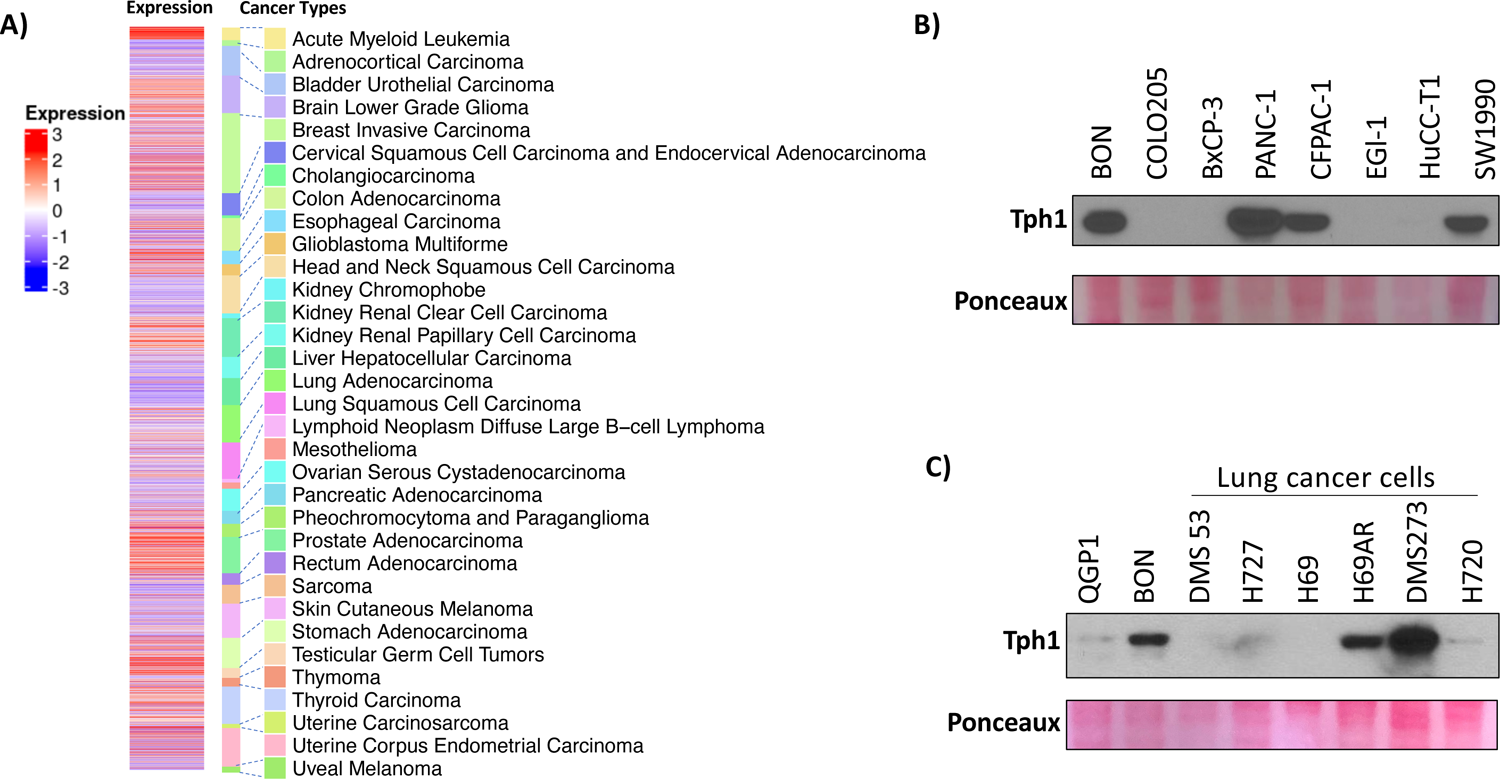
TPH1 expression in different cancer types and and cancer cell lines. (A) Expression levels of TPH1 gene across different cancer types according to TCGA database. (B) Protein levels of Tph1 in colon, pancreas, and cholangiocarcinoma cell lines. (C) Protein levels of Tph1 in lung cancer cell lines. Ponceaux staining of Western blots serves as loading controls.

### Inhibition of Tph1 does not affect BON cell viability or growth *in vitro*

BON cells synthesize large amounts of serotonin via Tph1, yet previous studies have reported no effect of the Tph1 inhibitor telotristat on their growth (Herrera-Martinez et al., 2020). A similar experiment to investigate the effect of TPH1 inhibition *in vitro* was accomplished by treating BON cells with 0.01-3 μM of telotristat and quantifying cell viability with the metabolic dye resazurin (Figure 3A). No significant change in cell viability was observed between control and treated cells at any concentration. To further investigate the effect of Tph1 inhibition, stable *TPH1* shRNA knockdown cell lines (BON shTPH1 #1 and #2) were generated and confirmed lower expression of Tph1 protein by western blot (Figure 3B). The growth rate of these cells over 6 days compared to BON cells expressing a control shRNA sequence (BON shControl) demonstrated no significant differences in knockdown cell lines versus control cells (Figure 3C).

**Figure 3.**
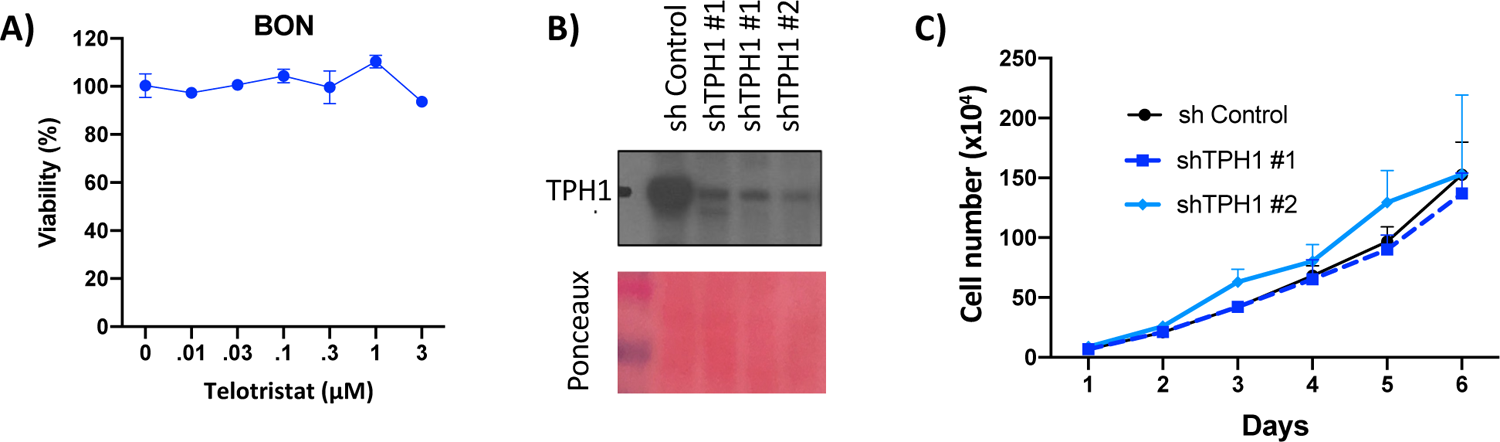
Effect of Tph1 inhibitor and *TPH1* knockdown in BON cells. (A) The effect of telotristat on BON cells in culture. (B) Clones of BON cells expressing shRNAs against *TPH1* (shTPH1#1 and shTPH1#2) showed decreased expression of Tph1 protein detected by western blot. Ponceaux stain depicts amount of protein loaded per lane. (C) *TPH1* knockdown cells divide at similar rates as control BON cells.

### BON *TPH1* knockdown cells form smaller tumors *in vivo*

Although Tph1 inhibition did not impair BON cell growth *in vitro*, the effect of these manipulations on *in vivo* growth was determined, as serotonin has been shown to support the growth of fibroblasts and endothelial cells, key components of the tumor microenvironment (TME) (Svejda et al., 2010, Blazevic et al., 2022). When 1 million BON parental, BON shControl, or BON shTPH1#1 and #2 cells were injected in the flank of NSG mice tumor volume data showed both BON shTPH1 cell lines form smaller tumors *in vivo* compared to control cells (Figure 4A & 4B) over 4.5 weeks. At the time of tumor harvest, the average weight of BON shTPH1 #1 and #2 knockdown tumors were approximately 0.1 g, significantly lower than the 0.3 g average weight of control tumors (Figure 4C). In addition, BON shControl tumors were dark red and appeared more vascularized in comparison to the light pink BON shTPH1 tumors (Figure 4D).

**Figure 4.**
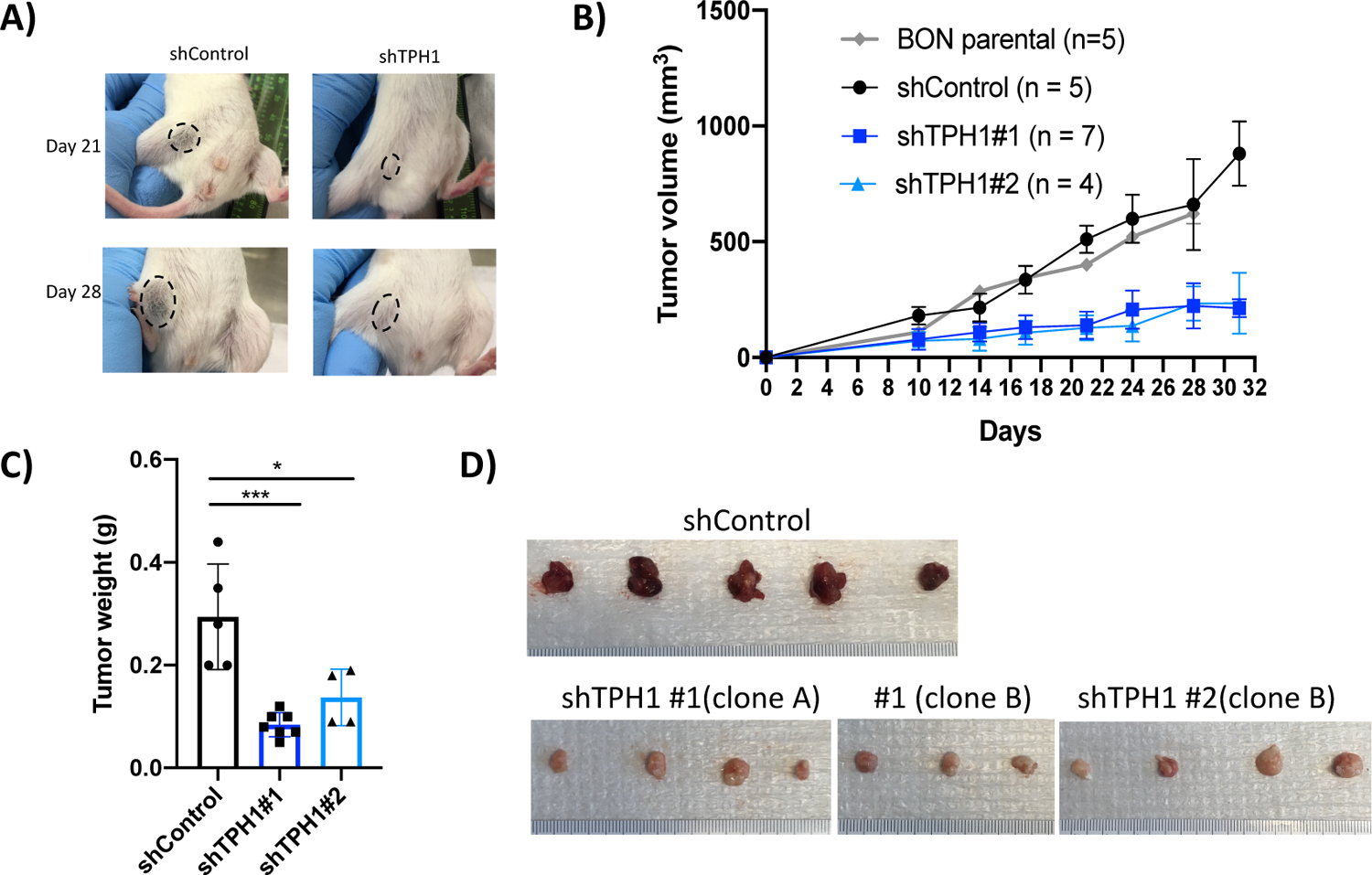
Characterization of BON xenografts with *TPH1* knockdown. (A) Xenograft model of BON cells expressing control shRNA (shControl) and shRNA targeting *TPH1* (shTPH1). (B) Tumor volume measurements with respect to time in days. (C) Weight of xenograft tumors from BON cells expressing shControl and 2 different shRNA targeting *TPH1* (shTPH1 #1 and shTPH1 #2) plotted as mean ± SD. P <0.05 was depicted with *. P <0.0001 was depicted with ****. (D) Images of BON xenograft tumors expressing the control shRNA or shRNAs targeting the *TPH1* gene.

### *TPH1* knockdown impairs tumor angiogenesis

The difference in tumor color from the xenograft models (Figure 4D) prompted investigation of the effect of *TPH1* knockdown on tumor endothelial cell density. IF staining for the endothelial cell marker CD31 on frozen xenograft tumor sections demonstrated less CD31 expression in the BON shTPH1 tumors compared to BON shControl tumors (Figure 5A). Vascular endothelial growth factors are key mediators of angiogenesis in the TME. *VEGF-A*, *VEGF-B*, *VEGF-C* gene expression in xenograft tumors detected by RT-qPCR showed expression of these genes were all reduced in shTPH1 xenograft tumors compared to shControl tumors, and this difference reached statistical significance for *VEGF-B* and *VEGF-C* (Figure 5B). Tube formation assay using human umbilical cord vein endothelial cells (HUVECs) cultured in conditioned media (CM) from BON shControl or BON shTPH1 cell lines with and without exogenous serotonin supplementation was assessed. After 2 days of incubation, bright field images of HUVECs were taken and tube formation based on tube length was quantified (Huuskes et al., 2019). Tube formation was decreased in HUVECs cultured in shTPH1 CM compared to shControl CM, but this effect could be reversed by exogenous serotonin supplementation (Figure 5C & 5D).

**Figure 5.**
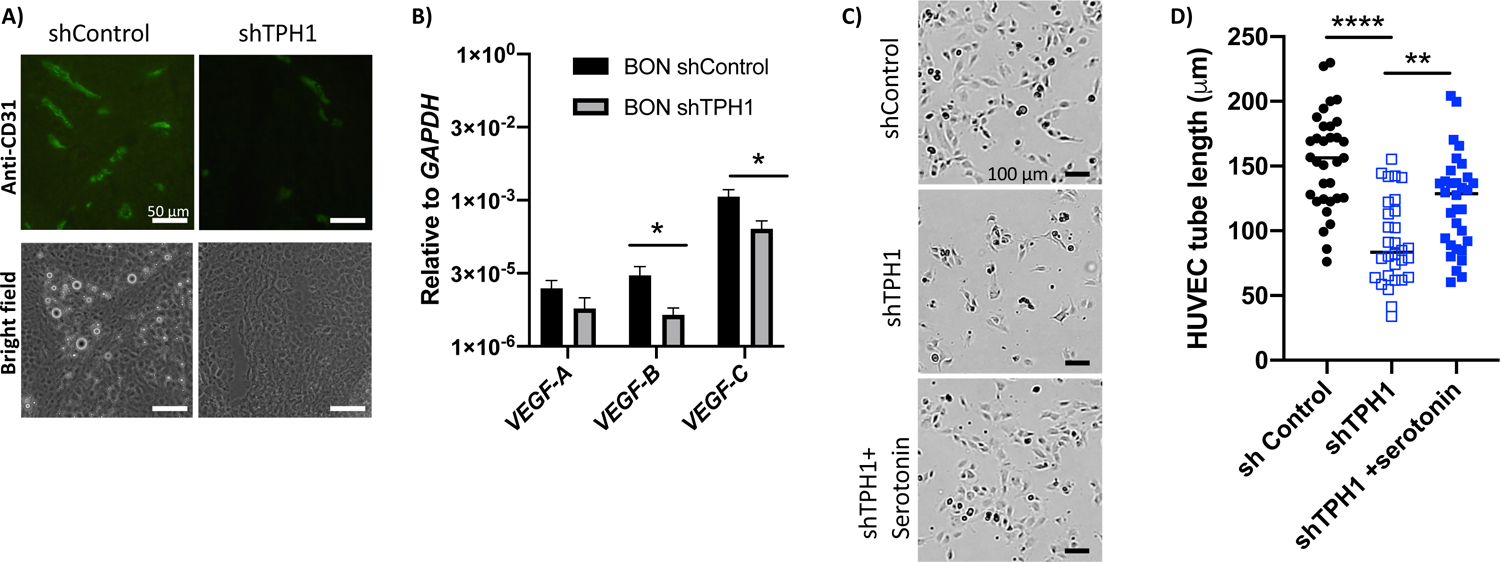
*TPH1* knockdown decreases angiogenesis. (A) Immuno-fluorescent staining for endothelial cells using antibody specific against CD31 in BON shControl tumors compared to shTPH1 tumors (n = 4 tumors/group). (B) Expression of vascular endothelial growth factors (*VEGF-A*, *VEGF-B*, *VEGF-C*) relative to *GAPDH* in BON tumors with shControl compared to shTPH1 plotted as mean ± SD. P <0.05 was depicted with *. (C) Tube formation assay with human umbilical cord vein endothelial cells (HUVECs) cultured in conditioned media from BON cells with shControl, shTPH1, or shTPH1 + 1 μM of serotonin. Shown are representative images from 3 independent experiments. (D) Quantification of HUVEC tube length from experiment described in C are represented as mean ± SD. P <0.01 was depicted with **. P <0.0001 was depicted with ****.

### Pharmacologic inhibition of Tph1 with telotristat reduces tumor growth *in vivo*

Given genetic knockdown of *TPH1* decreased tumor growth and vascularization in a mouse xenograft model (Figures 4 & 5), pharmacologic inhibition of Tph1 with the FDA-approved drug telotristat was tested on tumor growth. Implantation of 1 million BON cells in the flanks of 10 NSG mice was used to generate subcutaneous xenograft tumors. Ten-days post tumor cell injection, mice were randomized into control and telotristat treatment groups. Mice were given a vehicle control saline solution or 30 mg/kg telotristat by IP injections 5 times a week for 17 days. Tumor volume was measured twice per week, and telotristat treatment inhibited tumor growth (Figure 6A). At time of tumor harvest, the average weight of TE treated tumors was significantly less than control tumors (average of 0.15 g vs 0.30 g, P<0.001) (Figure 6B). Additionally, TE treated tumors appeared lighter in color and less vascularized compared to control tumors (Figure 6C). The effect of telotristat was tested on liver metastatic xenograft tumor models generated via intra-splenic injection of BON cells expressing a firefly luciferase reporter gene (Kaemmer et al., 2021) in NSG mice. Seven days-post injection, mice were randomized into treatment groups and given a vehicle control saline solution or 30 mg/kg telotristat by IP injections 5 times a week for the next 14 days. Tumor burden was quantified by *in vivo* bioluminescent imaging at day 7 and day 21 post-tumor cell injection. At day 21, decreased luminescent signal was observed from mice that received telotristat treatment compared to the control group, indicating telotristat reduced metastatic tumor burden (Figure 6D & 6E).

**Figure 6.**
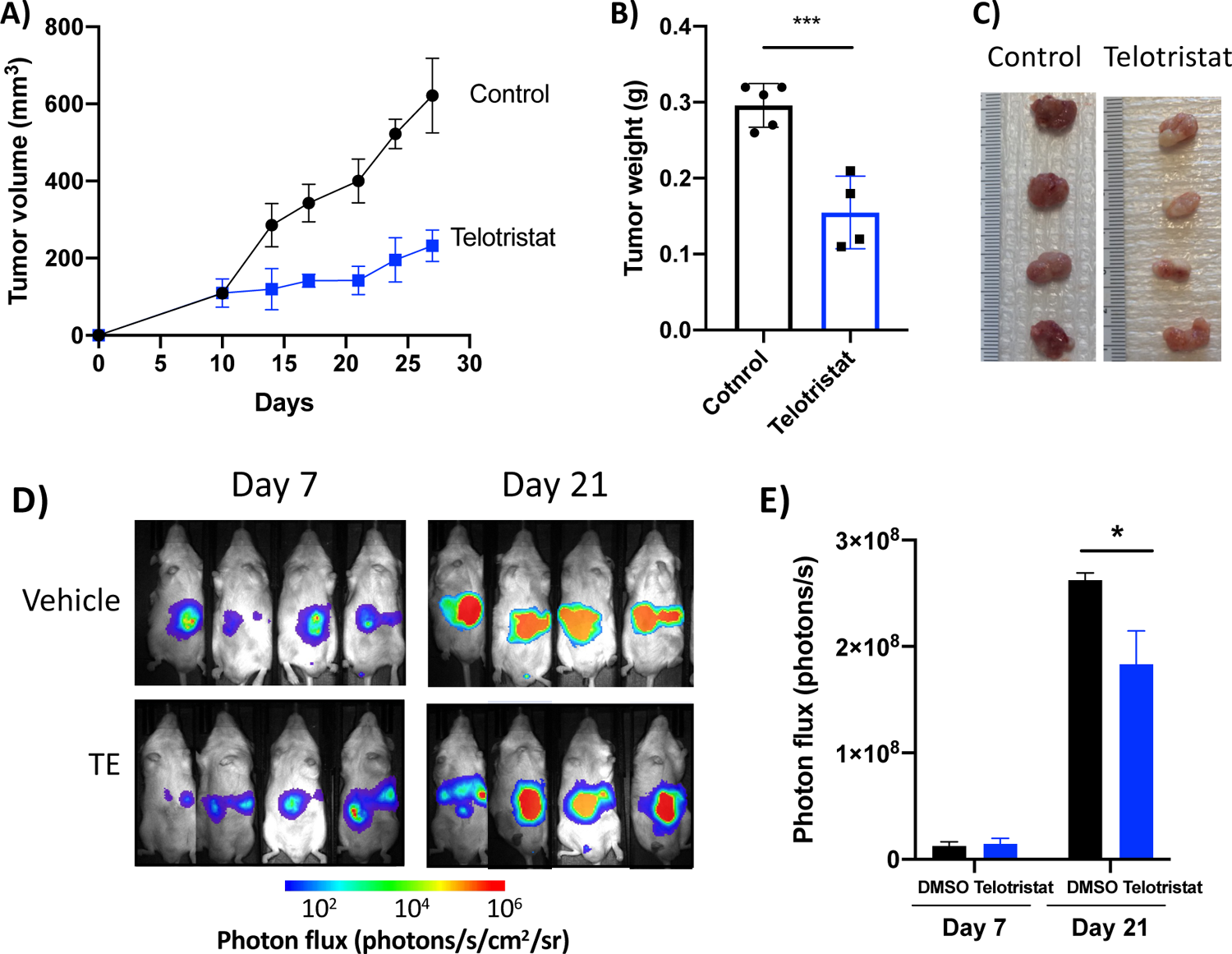
Inhibition of TPH1 with telotristat reduces tumor growth *in vivo*. (A) Tumor volume measurement with respect to time. Ten-day post BON cell tumor implantation, 5 mice were given a vehicle control and 4 mice were given telotristat 5 times per week at 30mg/Kg. (B) Weight of xenograft tumors from BON cells with and without telotristat treatment plotted as mean ± SD. P <0.001 was depicted with ***. (C) Picture of BON xenograft tumors with and without telotristat treatment. (D) Liver metastases model of BON xenografts established by intrasplenic injections of luminescent BON cells in NSG mice. Tumor burden was monitored at day 7 and day 21 post-tumor cell injection for vehicle control and telotristat treated mice. Photon flux is represented as photons/s/cm^2^/sr. (E) Quantification of bioluminescent signal as total photon flux (photons per second) are represented as mean ± SD. P <0.05 was depicted with *.

## Discussion

Tph1 is the rate limiting enzyme for the synthesis of serotonin, a biogenic amine with autocrine and paracrine functions in regulating tumor growth. Several studies have demonstrated a link between Tph1 and serotonin in cancer development. However, the role of Tph1 and serotonin biosynthesis in NENs remains unclear. Although NEN cancer cells express Tph1 (Figure 1&2) and have 5-HT2B receptors on their cell surface, we found no effect of Tph1 inhibition using telotristat or *TPH1* knockdown on the growth of BON cells *in vitro* (Figure 3). One reason for this could be related to how the BON cells were cultured. This study and another (Herrera-Martinez et al., 2020), BON cells were cultured in a rich medium supplemented with excess glutamine and 10% fetal bovine serum (FBS). It has been reported that glutamine can stabilize *TPH1* mRNA levels (Park et al., 2023) thereby leading to greater abundance of the Tph1 enzyme, reducing the effect of telotristat in these experiments. Interestingly, Drozdov et al. used a suicide inhibitor of Tph1 (7-hydroxytryptamine) and reported decreased ERK and JUN signaling pathways in lung NEN cells (H720, H727) and EBV-transformed lymphoblastoid cells (KRJ-I, originally misidentified as SBNET) (Drozdov et al., 2009). High levels of FBS in the culture medium or high levels of growth factor production by BON cells could saturate the tumor cell proliferation signal to constitutively activate the MAPK signaling pathways to prevent telotristat, a competitive TPH1 inhibitor, from inhibiting cell growth *in vitro*.

In distinct contrast to results from cultured cell studies, our data revealed for the first time that inactivation of Tph1, using both genetic and pharmacological inhibition, significantly suppresses NEN tumor growth *in vivo* (Figures 4 & 6). One likely mechanism for this reduced growth is a reduction in serotonin signaling that promotes angiogenesis. Endothelial cells express multiple 5-HT receptor subtypes and serotonin at low concentrations can activate pro-growth signaling cascades in endothelial cells (Zamani and Qu, 2012). Our data also showed BON *TPH1* knockdown tumors to be less vascularized than control tumors (Figure 4D), indicating that serotonin is an important regulator of angiogenesis in these tumors. Less endothelial cell staining and decreased *VEGF-B* and *VEGF-C* expression in *TPH1* knockdown tumors (Figures 5A & B) was also observed. Notably, addition of serotonin to conditioned medium collected from BON *TPH1* knockdown cells was able to restore endothelial cell survival and tube formation (Figures 5C & D), supporting the serotonin dependence of the impaired angiogenic phenotype caused by *TPH1* knockdown (Figures 5C & D).

Recent studies have revealed important functions of serotonin regulation in tumorigenesis and drug resistance in in PDAC (Chaudhary et al., 2021, Jiang et al., 2017), colon cancer (Liu et al., 2021, Schneider et al., 2021), breast cancer (Gautam et al., 2016), and others. In our analysis of Tph1 expression in small cell lung carcinoma cells, high Tph1 was detected in the doxorubicin resistant H69AR cell line as compared to the parental H69 cell line which is sensitive to doxorubicin (Figure 2C). In addition, high Tph1 expression was also observed in the DMS 273 cell line (Figure 2C) which is derived from a patient tumor resistant to chemotherapy and radiotherapy. This suggests that high Tph1 levels could contribute to acquired drug resistance in tumors. Mechanisms by which serotonin might activate drug resistance pathways have not been reported, however, inhibiting Tph1 in combination with other anti-cancer therapies is a strategy currently under exploration. In preclinical studies, telotristat treatment enhanced the efficacy of temozolomide in a glioma xenograft model (Zhang et al., 2022). Similarly, Tph1 inhibition combined with gemcitabine further decreased tumor growth relative to gemcitabine alone in PDAC xenograft tumors (Chaudhary et al., 2021). Those results support an ongoing clinical trial in PDAC (NCT03910387) to determine the effect of telotristat ethyl at promoting weight stability in patients treated with gemcitabine and paclitaxel. A secondary objective of the trial is to determine if the telotristat ethyl combination with gemcitabine and paclitaxel has improved anti-tumor effects. Currently, another clinical trial that is actively enrolling NET patients seeks to determine if inhibition of serotonin production using telotristat ethyl alone has a cytostatic effect on NETs and/or if it enhances the antitumor activity of peptide receptor radionuclide therapy using ^177^Lutetium-Dotatate (NCT04543955).

In conclusion, few effective medical therapies are available for treating NENs. Targeting TPH1 activity to suppress tumor growth has not been adequately explored in NENs but is highly relevant given the excessive production of serotonin by some types of NENs and its approval in NEN patients with carcinoid syndrome. Our study is the first to demonstrate that Tph1 inhibition leads to decreased NEN tumor growth *in vivo*. Accumulating evidence in the literature suggest combination therapy targeting Tph1 together with other anti-cancer agents would be a promising approach since telotristat ethyl therapy is associated with few negative side effects in patients (Kulke et al., 2014, Pavel et al., 2018). Along with emerging studies reporting the importance of serotonin regulation in NENs (Soldevilla et al., 2021), inhibiting this enzyme could be a safe and impactful strategy for targeting the metabolic vulnerability of these highly drug-resistant cancers.

## Declaration of interest

C. Chandrasekharan was an employee/paid consultant for Lexicon. C.H.F. Chan received research support from Angiodynamics, Checkmate Pharmaceuticals, Optimum Pharmaceutics, and Regeneron for unrelated research projects. Other authors have no conflicts of interest to disclosed.

## Funding

This work was supported by the University of Iowa NET SPORE P50CA174521, CA260200, CA217797, CA086862, NANETS/NETRF BTSI award and Holden Comprehensive Cancer Center GI-Cancer Pilot fund.

## Contributions

PHE, DHT, JRH, CHC, JD, CC, MH, DRS, and DEQ conceived the study, wrote and edited this manuscript. PHE, DHT, CGT, LCB, MR, GL, CAK, EA, CHC, AMB performed experiments and analyzed data.

## Acknowledgements

We thank Dr. Thomas O’Dorisio for guidance and support at the initial phase of this project, Drs. M.B. Evers and C.M. Townsend for BON cells, and Dr. L.R. Roberts for HuCC-T1 and EGl-1 lines.

